# Brain-wide connectivity patterns of feedforward and feedback cortico-cortical neurons in the mouse secondary visual cortex

**DOI:** 10.1101/2025.07.08.663552

**Authors:** Richard G. Dickson, Matthew W. Jacobs, John M. Ratliff, Alec L.R. Soronow, Faye An, Walid A. Yuqob, Euiseok J. Kim

## Abstract

Feedforward and feedback cortico-cortical neurons are distinct yet spatially intermingled subtypes distributed across cortical layers, playing specialized roles in sensory and cognitive processing. However, whether their presynaptic inputs differ to support these functions remains unknown. Using projection-and layer-specific monosynaptic rabies tracing, we mapped brain-wide long-distance inputs to multiple feedforward and feedback neuron types in VISl (also known as LM), the mouse secondary visual cortex. Overall, long-distance input patterns for these feedforward and feedback neurons were largely similar, as all received the majority of their inputs from VISp, the primary visual cortex, along with substantial inputs from various other cortical and visual thalamic regions. Despite their similarities, these feedforward and feedback types differed in the proportion of long-distance cortical inputs originating from specific visual, retrosplenial, and auditory cortices. These findings reveal the input connectivity patterns of cortico-cortical neurons based on feedforward and feedback projections, providing an anatomical framework for future studies on their functions and circuit integration.

**KEY POINTS:**

1) Feedforward and feedback cortico-cortical projection neurons in VISl form largely distinct populations that are distributed across multiple cortical layers.
2) Four different feedforward and feedback cortico-cortical projection neuron types receive broadly similar patterns of long-distance input across the brain.
3) Despite overall similarities, feedforward and feedback neurons differ in the relative proportions of inputs from specific areas, including visual, retrosplenial, and auditory cortices.

## INTRODUCTION

The mammalian cortex is responsible for high-level perception and cognition and is composed of numerous interconnected cortical areas (Felleman and Van Essen, 1991; Oh et al., 2014; Gămănuţ et al., 2018). These areas communicate through long-distance inter-areal connections mediated by excitatory cortico-cortical projection neurons (CCPNs). Two key organizational features of the mammalian cortex are its six-layered laminar structure and its hierarchical organization (Felleman and Van Essen, 1991; Cajal, 1995). Each cortical layer is considered as a different information-processing unit (Harris and Shepherd, 2015). CCPNs in a specific layer of each cortical area can be further divided into feedforward projection neurons, which send axons to higher-order cortical areas, and feedback projection neurons, which project to lower-order cortical areas. These two neuronal types are thought to process distinct types of information, feedforward neurons primarily transmitting sensory input from the external world and feedback neurons modulating contextual factors such as attention, brain states, and background information (Hupé et al., 1998; Nassi et al., 2013; Keller et al., 2020; Debes and Dragoi, 2023). Despite their distinct roles, these feedforward and feedback neurons are spatially intermingled within cortical layers (Berezovskii et al., 2011). Recent advances have elucidated their functional roles (Keller et al., 2020; Fişek et al., 2023), however, it remains unclear whether feedforward and feedback projection neurons within a specific layer of a given cortical area receive distinct or similar brain-wide presynaptic inputs. Uncovering this would provide insight into their proposed functional differences.

The mouse visual cortex serves as an ideal model for investigating the brain-wide connectivity of layer-and projection-specific cell types at a cellular level. This is due to the availability of viral genetic tools that enable precise labeling of neurons based on their axonal projections (e.g., retrograde tracer such as AAVretro) and laminar identity (e.g., layer-specific Cre-expressing transgenic mouse lines) (Gerfen et al., 2013; Tervo et al., 2016; Chatterjee et al., 2018; Matho et al., 2021). Additionally, the hierarchical organization of mouse visual cortical areas has been well characterized both anatomically and functionally (Harris et al., 2019; Siegle et al., 2021; D’Souza et al., 2022). The mouse visual cortex comprises the primary visual cortex (VISp) and at least ten different higher visual areas (HVAs), including the lateromedial (VISl) and posteromedial (VISpm) visual cortices (Wang and Burkhalter, 2007). In the cortical hierarchy, VISp represents the lowest level, VISl occupies an intermediate position, and VISpm is at a higher level (Harris et al., 2019; D’Souza et al., 2022). Within this framework, CCPNs in VISl that project to VISpm are defined as feedforward neurons, whereas those projecting to VISp are defined as feedback neurons (D’Souza et al., 2022). This makes VISl particularly suitable for studying spatially intermingled feedforward and feedback CCPNs.

Previous studies have mapped the monosynaptic inputs of layer-specific excitatory neuronal types (Vélez-Fort et al., 2014; DeNardo et al., 2015; Kim et al., 2015, 2020; Yao et al., 2023a). These studies suggest that long-distance input connectivity to specific layer-defined neuronal classes can be either distinct or largely comparable and that local connectivity varies (DeNardo et al., 2015; Yao et al., 2023a). However, it remains unclear whether feedforward and feedback CCPNs within the same cortical layer receive distinct brain-wide monosynaptic inputs, highlighting a critical gap in our understanding of cortical circuit organization.

To investigate the brain-wide monosynaptic input connectivity of layer-specific feedforward and feedback CCPNs in the mouse visual cortex, we employed the intersectional G-deleted rabies tracing method - cTRIO (cell-type specific inputs and outputs relationship) (Schwarz et al., 2015). Using an intersectional approach that combined layer-specific Cre recombinase with projection-specific tTA trans-activator expression, we defined distinct subtypes of feedforward and feedback CCPNs in layer 2/3 (L2/3) and 5 (L5) of VISl. We then mapped their long-distance monosynaptic inputs using rabies tracing, leveraging two avian receptors, TVA and TVA^66T^, for avian sarcoma leukosis virus glycoprotein EnvA (Wickersham et al., 2007; Miyamichi et al., 2013). Thus we are able to trace the monosynaptic inputs to neuronal subtypes which were previously inaccessible to brainwide mapping efforts. Our findings reveal that while the overall input patterns to L2/3 and L5 feedforward and feedback CCPNs in VISl are largely comparable, there are notable differences in the proportions of long-distance inputs from the visual, auditory, and retrosplenial cortices. The experimental design presented here provides a framework for investigating hierarchical brain-wide networks across different species and brain structures beyond the mouse visual cortex. This study also provides a comprehensive presynaptic input connectivity map that will serve as a foundation for future experimental and theoretical investigations into cortico-cortical neuronal functions.

## MATERIALS AND METHODS

### Resource and materials availability

Further information and requests for resources and materials should be directed to and will be fulfilled by the lead contact, Euiseok J. Kim. Microscopy and any additional data reported in this paper will be shared by the lead contact upon reasonable request.

### Experimental animals

GENSAT BAC transgenic *SepW1-Cre* NP39 and *Tlx3-Cre* PL56 mouse lines have been previously described (Gerfen et al., 2013). The transgenic mice were maintained on C57BL/6J backgrounds. C57BL/6J mice were used as wild-type. Both male and female mice were used. All mice were housed with a 12-hour light and 12-hour dark cycle and *ad libitum* access to food and water. All animal procedures were performed in accordance with the University of California, Santa Cruz animal care and use committee (IACUC)’s regulations.

### Virus preparation

All adeno-associated viruses (AAVs) and SAD B19 EnvA-pseudotyped glycoprotein (G)-deleted rabies virus (EnvA+RVdG) were produced by the Salk Viral Core GT3: scAAVretro-hSyn-BFP (1.26X10^12^ GC/ml), scAAVretro-hSyn-H2B-eGFP (1.16X10^13^ GC/ml), scAAVretro-hSyn-H2B-mCherry (1.21X10^13^ GC/ml), AAVretro-nEF-lox66/71-tTA (4.8X10^13^ GC/ml), AAV8-TRE-DIO-oG (5.92X10^12^ GC/ml), AAV8-TRE-DIO-eGFP-T2A-TVA (7.00X10^13^ GC/ml), AAV8-TRE-DIO-TVA^66T^eGFP-P2A-oG (7.40X10^13^ GC/ml), and EnvA+RVdG-mCherry (7.71X10^8^-1.43X10^10^) Infectious Unit (IU)/ml).

### Animal surgery for virus injection

For all surgeries anesthesia induction was with 100 mg/kg of ketamine and 10 mg/kg of xylazine cocktail via intraperitoneal injections, and anesthesia was maintained as needed with ∼1% isoflurane. Mice were mounted in a stereotax (RWD instruments) for surgery and stereotaxic injections. For rabies tracing experiments, *SepW1-Cre* NP39 or *Tlx3-Cre* PL56 mice received AAV helper injections at postnatal day (P)63-P125. 50-200 nl of AAVretro-nEF-lox66/71-tTA was injected into the center of the medial secondary visual cortex (VISpm), primary visual cortex (VISp), or anterior cingulate area (ACA) using the following coordinates: 1.6 mm rostral, 1.5 mm lateral relative to lambda and 0.4-0.6 mm ventral from the pia for VISpm or 1.1 mm rostral, 2.6 mm lateral to lambda and 0.45-0.67 mm ventral from the pia for VISp, 0.4 mm rostral, 0.4 mm lateral to bregma and 0.7-0.8 mm ventral from the pia for ACA. A 50 nl mixture of 9:1 AAV8-TRE-DIO-oG and AAV8-TRE-DIO-eGFP-T2A-TVA, a 50 nl mixture of 2:7:1 AAV8-TRE-DIO-oG, 1x PBS and AAV8-TRE-DIO-eGFP-T2A-TVA, or 50 nl of AAV8-TRE-DIO-TVA^66T^eGFP-P2A-oG was injected into the center of the lateral secondary visual cortex (VISl), using the following coordinates: 0.7 mm rostral, 3.65 mm lateral relative to lambda and 0.4-0.6 mm ventral from the pia. We injected AAVs using air pressure using a 1 ml syringe with 18G tubing adaptor and tubing. To prevent virus backflow, the pipette was left in the brain for 5-10 minutes after the completion of the injection. After recovery, mice were given water with ibuprofen (30 mg/kg) and subcutaneous carprofen (5 mg/kg). Two weeks after AAV helper injection, 100 nl of EnvA+RVdG-mCherry was injected into the same site in VISl using a 1 ml syringe-mediated air pressure. After recovery, mice were again given water with ibuprofen (30 mg/kg) and subcutaneous carprofen (5 mg/kg) and housed for seven days to allow for trans-synaptic rabies spread and fluorescent protein expression.

For retrograde colocalization studies, mice were prepared and head-fixed as previously with the same coordinates for VISpm, VISp, and ACA as above. For triple AAV colocalization experiments, combination of 50-100 nl of scAAVretro-hSyn-BFP, scAAVretro-hSyn-H2B-mCherry, scAAVretro-hSyn-H2B-eGFP were injected into one of VISp, VISpm, and ACA. AAV was left to express for 18-21 days before the tissue harvest and histology. For VISpm/VISp/ACA chemical tracer colocalization experiments, 50-100 nl of cholera toxin subunit B (CTB) conjugated to Alexa fluor 647 (CTB-647, 1 mg/ml, Invitrogen C34778), Alexa fluor 555 (CTB-555, 1 mg/ml, Invitrogen C22843) and/or Alexa fluor 488 (CTB-488, 1 mg/ml, Invitrogen C34775) was injected into VISpm, VISp, and/or ACA. Three to six days later, the tissue was harvested for histology. Different fluorophore colors were rotated between the three injection sites across different animals. Detailed information of injections on individual mice can be found in Table 1.

**Table 1.**
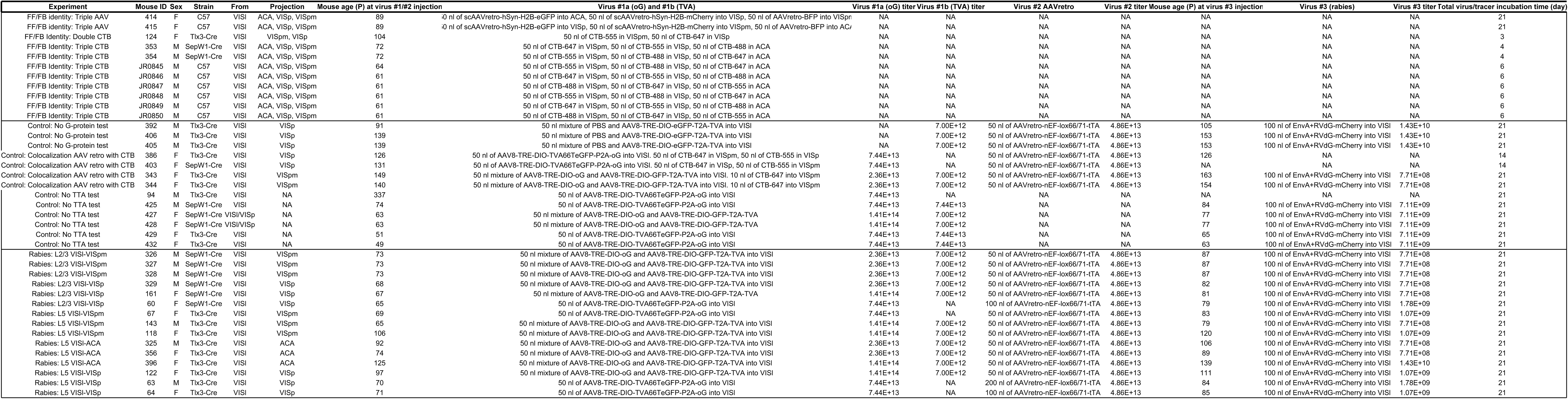
Animal and experimental condition summary.

### Histology and image analysis

Brains were harvested after trans-cardiac perfusion using phosphate-buffered saline (PBS) followed by 4% paraformaldehyde (PFA). Brains were dissected out from skulls and post-fixed with 2% PFA and 15% sucrose in PBS at 4°C overnight and then immersed in 30% sucrose in PBS at 4°C before sectioning. Using a freezing microtome, 50-100 µm coronal brain sections were cut and stored in cryogen (30% ethylene glycol, 30% glycerol, 30% ddH_2_0, 10% 10x PBS) at 4°C, every other section was stained and mounted for data analysis and quantification. Before staining, sections were washed 3x 10 minutes in PBS and permeabilized and blocked in PBS/5% normal donkey serum (Millipore S30-100ML)/1% Triton-X 100 (Sigma-Aldrich X100-100ML) for two hours at room temperature. To enhance eGFP and mCherry signals, free-floating sections were incubated at 4°C overnight with goat anti-GFP (1:1000; Rockland 600-101-215, RRID:AB_218182) and rabbit anti-dsRed (1:500; Clontech 632496, RRID:AB_10013483) primary antibodies in PBS/0.5% normal donkey serum/0.1% Triton-X 100, followed by 3x 10 minutes in PBS wash and incubation with the appropriate secondary antibodies conjugated with Alexa 488 or 568 (1:500; Invitrogen A-11055 (RRID:AB_2534102) or A-10042 (RRID:AB_2534017), respectively). Sections were counterstained with 10 μM DAPI (4’,6-diamidino-2-phenylindole) in PBS for 30 minutes to visualize cell nuclei, followed by 3x 10 minutes in PBS wash. Immunostained tissue sections were mounted on slides with a polyvinyl alcohol mounting medium containing DABCO (PVA-DABCO) and allowed to air-dry overnight.

All sections were scanned with a 10x/0.45 NA (wd-2.0 mm) objective on a Zeiss AxioImager Z2 Widefield Microscope using tile scan and Z-stack (every 6-8 µm). Representative images were taken with the Zeiss 880 Confocal Microscope with 10x/0.45 NA (wd-2.0 mm) or 40x/ 0.95 NA corr (wd-0.25 mm) objectives. Scanned images were first processed in Zeiss ZEN 3.11 with DAPI images for alignment and fluorescent images for eGFP+ or mCherry+ cell detections exported for processing. Subsequent image files were processed and analyzed using Bell Jar for long-distance input mapping (Soronow et al., 2025). For alignment, the center Z-stack DAPI images were used to predict the section’s xy coordinates and cut angle for alignment to the Allen Brain Common Coordinate Framework reference atlas (Wang et al., 2020). Alignments were then fine-tuned in Bell Jar to ensure fit and minimize counting rabies-labeled cells resulting from minor TVA expression leakage in non-Cre-expressing neurons (Wall et al., 2010). Layers for VISpm and VISp were also manually fine-tuned to ensure accuracy. The exported Zeiss ZEN files with mCherry+ signals were used for automated cell counting. Using Bell Jar, max projection images were produced from the Z-stacks, and these max projections were then sharpened in the program before detection. Detection thresholds were kept stable unless manual counting validation showed less than 90% accuracy of cell labeling, in which case the confidence interval was adjusted (three out of 15 brains) to ensure accurate counting. Cell detection and brain alignment were then integrated, and the long-distance ipsi-hemispheric cortical input fractions were calculated and graphed using GraphPad Prism 10.1.1. For thalamic input proportions, areas and cells were drawn and counted manually on Zeiss ZEN 3.11 using the Allen Brain Common Coordinate Framework reference atlas (Wang et al., 2020).

For VISpm/VISp/ACA CTB colocalization experiments, all CTB labeled cells were counted manually due to weaker fluorescent signals that could not reliably be detected using Bell Jar’s automated cell detection. For each animal, three sections corresponding to the same three VISl containing plates from the Allen CCFv3 were selected for counting. Using Fiji, VISl and laminar borders were drawn over DAPI images, cells were labelled individually and then overlayed labels were checked for co-localization.

### Quantification and Statistical analysis

The values of n and what n represents are reported in results and figure legends. For statistical significance, two-way ANOVA with Tukey’s multiple comparisons test was performed with GraphPad Prism 10.1.1. A one-way ANOVA with the Fisher’s Least Significant Difference (LSD) post hoc for multiple comparisons test was performed using IBM SPSS 30. Unless otherwise noted, statistical significance is indicated as follows: *p < 0.05; **p < 0.01; ***p < 0.001; ****p < 0.0001. In Fig. 4C, significance is denoted differently: *(black) p < 0.05 and *(magenta) p < 0.01.

## RESULTS

### Feedforward and feedback cortico-cortical projection neurons of VISl are largely non-overlapping and distributed across layers

We first investigated the long-distance projection targets of CCPNs in VISl using published data from the Allen Mouse Brain Connectivity Atlas (Harris et al., 2019). This dataset provides high-resolution maps of eGFP+ axonal projections in adult mouse brains, generated by injecting AAV1-CAG-FLEx-eGFP into Cre-expressing transgenic mice to label specific neuronal populations in different brain regions. To examine CCPNs in VISl, we analyzed data from *Cux2-IRES-Cre* and *Tlx3-Cre* mice, which label layers 2-4 (L2-4) and L5 CCPNs, respectively. Whole-brain projection maps revealed that both L2-4 and L5 CCPNs extend axons to more than 13 cortical areas, including visual areas as VISp, VISpm, and VISpor and non-visual areas such as the temporal association cortex (TEa) and the anterior cingulate area, dorsal part (ACAd) (Fig. 1A-B). Quantitative analysis of the axonal projection volumes across these regions showed that L2-4 and L5 CCPNs exhibit similar projection patterns, with the highest projection volume in VISp (39.43 ± 3.20% in *Cux2-IRES-Cre* and 29.75 ± 3.80% in *Tlx3-Cre*, mean ± SEM), followed by VISpm, VISpor, VISli, and VISrl (Fig. 1B). Notably, L2-4 CCPNs exhibit sparse axonal projections to ACAs (1.10 ± 0.28%), whereas 3.55 ± 0.95% of the total axonal projections from L5 CCPNs target ACAd (Fig. 1B).

**Figure 1.**
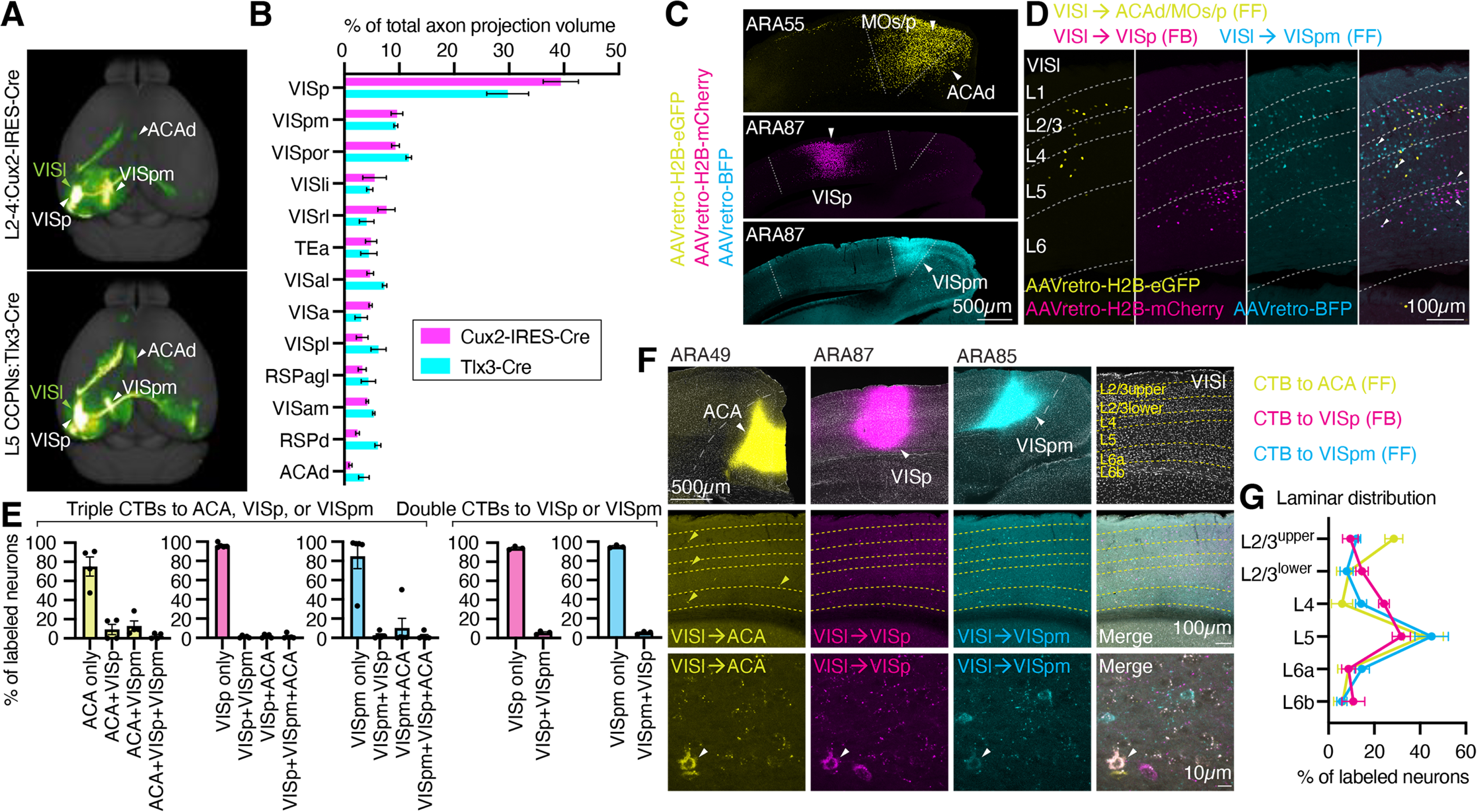
Whole-brain axonal projections and largely segregated feedforward and feedback CCPNs in VISl. **(A)** Top-down views of whole-brain 3D reconstructions of axonal projections from VISl. (Top) L2-4 CCPNs in a *Cux2-IRES-Cre* brain (Experiment: 287495026). (Bottom) L5 CCPNs in a *Tlx3-Cre* brain (Experiment: 520750912). Data adapted from the Allen Brain Connectivity Atlas (https://connectivity.brain-map.org/projection). **(B)** Percentage of total axon projection volume across 13 cortical areas from L2-4 VISl CCPNs in *Cux2-IRES-Cre* (magenta, n = 2 mice) and L5 VISl CCPNs in *Tlx3-Cre* (cyan, n = 3 mice). **(C)** Confocal microscope images showing the injection sites of AAVretro-H2B-eGFP, AAVretro-H2B-mCherry, and AAVretro-BFP targeting ACAd/MO, VISp, and VISpm, respectively. Allen Reference Atlas (ARA) section numbers are in the top left corner. **(D)** Representative confocal microscope images showing VISl CCPNs retrogradely labeled based on their axonal projections to ACAd/MO, VISp, or VISpm. Arrows indicate neurons that project to more than one of these target areas. **(E)** Proportions of VISl neurons projecting to multiple target areas. Triple CTB injection experiments show the fraction of VISl neurons projecting to single, double, or all three target areas (ACA, VISp, VISpm). Counts per CCPN type across all animals were: 94 cells for VISl→ACA, 523 cells for VISl→VISp, and 573 cells for VISl→VISpm (n =4 mice). Double CTB injections show the fractions of VISl neurons projecting to one or both target areas (VISp or VISpm). Counts per CCPN type across all animals were: 601 cells for VISl→VISp and 763 cells for VISl→VISpm (n = 3 mice). **(F)** Representative confocal images of CTB-labeled VISl neurons. Top row: Injection sites of CTB tracers conjugated to three fluorophores targeting ACA (yellow), VISp (magenta), and VISpm (cyan); ARA section numbers are indicated above each panel. Middle and bottom rows: VISl CCPNs retrogradely labeled according to their projections. Yellow arrows highlight neurons projecting to ACA, and white arrows indicate neurons projecting to ACA, VISp, and VISpm. **(G)** Percentage distribution of the somata of three VISl CCPN types across cortical layers. A total of 1190 retrograde labeled neurons were analyzed in 6 mice. Data are represented as mean ± SEM.

Next, we investigated whether CCPNs in VISl that project to three major cortical target areas (VISp, ACA/MOs (secondary motor cortex), and VISpm) represent distinct or overlapping cell types at the single-cell level. To retrogradely label these projection neurons with three different fluorescent markers, we stereotaxically injected AAVretro-hSyn-H2B-mCherry into VISp, AAVretro-hSyn-BFP into ACA/MOs, and AAVretro-hSyn-H2B-eGFP into VISpm in wild-type adult C57BL/6J mice (Fig. 1C). Thus, VISl neurons projecting to VISpm, VISp, and ACA/MOs, designated as VISl→VISpm, VISl→VISp, and VISl→ACA, respectively. In VISl of these mice, labeled CCPNs were distributed across layers 2-6. The majority of neurons were labeled by a single fluorescent protein (VISl→VISpm: 90.97 ± 2.08%; VISl→VISp: 92.25 ± 5.04%; VISl→ACA: 87.94 ± 4.60%, n = 2 mice) with relatively few exhibiting double or triple labeling (Fig. 1C). These findings indicate that CCPNs projecting to VISpm or ACA/MOs (feedforward, FF) and VISp (feedback, FB) do not extensively share axonal collaterals consistent with a previous study (Berezovskii et al., 2011).

To evaluate whether this observation is consistent when using chemical tracers that avoid potential viral co-transduction effects of AAVretros, we stereotaxically injected 50 nl of cholera toxin subunit B (CTB) conjugated to Alexa Fluor 647, 555, or 488 into the ACA, VISp, and VISpm, respectively, alternating fluorophores across animals. In VISl of these triple CTB injected mice, most neurons were labeled by a single fluorescent CTB (VISl→ACA: 75.11 ± 10.04%; VISl→VISp: 95.93 ± 1.06%; VISl→VISpm: 85.00 ± 12.92%, n= 4 mice), with relatively few neurons exhibiting double or triple labeling (Fig. 1E, F). In a separate set of experiments, we injected 100 nl of CTB conjugated to Alexa Fluor 647 or 555 into VISp or VISpm, respectively, to maximize labeling of neurons projecting to these areas while minimizing unintended spread to adjacent regions. Even under these double injection conditions, the majority of VISl neurons were labeled by a single fluorescent CTB (VISl→VISp: 94.40 ± 1.05%; VISl→VISpm: 94.95 ± 0.87%, n=3 mice) (Fig. 1E). Therefore, our results indicate that VISl→VISpm, VISl→VISp, and VISl→ACA CCPNs are largely distinct neuronal subpopulations.

CCPNs are distributed across all cortical layers from L2/3 to L6. To determine the laminar distribution of the three VISl CCPN types projecting to ACA, VISp, and VISpm, we quantified their soma locations as percentages across layers (Fig. 1G). All three types were present throughout L2-6, with the highest proportions in L5 (VISl→ACA: 41.50 ± 8.62%; VISl→VISp: 31.82 ± 3.97%; VISl→VISpm: 45.07 ± 7.29%; n = 3 mice, Fig. 1G). Notably, VISl→ACA CCPNs were more abundant in the upper division of L2/3 (L2/3^upper^) compared to the lower division (L2/3^lower^) (28.56 ± 3.91% vs. 9.08 ± 5.48%, Fig. 1G). VISl→VISpm CCPNs showed a similar trend, with higher proportions in L2/3^upper^ than L2/3^lower^ (12.29 ± 1.77% vs. 7.93 ± 2.87%, Fig. 1G). In contrast, VISl→VISp CCPNs were more abundant in L2/3^lower^ compared to L2/3^upper^ (14.66 ± 2.67% vs. 9.58 ± 3.40%, Fig. 1G). Despite these laminar differences, all three CCPN types displayed substantial overlap across the full depth of L2/3 and within L5.

### Brain-wide monosynaptic input mapping of feedforward and feedback CCPNs in VISl

We examined whether long-distance inputs to feedforward and feedback neurons align with one of three possible organizational models (Fig. 2A). In Model 1, these neurons receive inputs from the same areas they project to, establishing a reciprocal connectivity pattern that facilitates looped computations. In Model 2, feedforward and feedback neurons differ in the type of information they relay; feedforward neurons preferentially receive inputs from lower sensory areas such as VISp and the dorsal lateral geniculate nucleus of the thalamus (LGd), while feedback neurons receive inputs from higher cortical areas that convey contextual information such as higher cortical areas and the lateral posterior thalamic nucleus (LP). In Model 3, both feedforward and feedback neurons receive inputs from the same brain regions, making them anatomically indistinguishable. In this scenario, despite similar long-distance anatomical inputs, feedforward and feedback neurons may exhibit distinct functional roles due to differences in local circuit computations. Alternatively, the long-distance inputs themselves may differ functionally, even if their anatomical connectivity proportion from each brain region appears similar.

**Figure 2.**
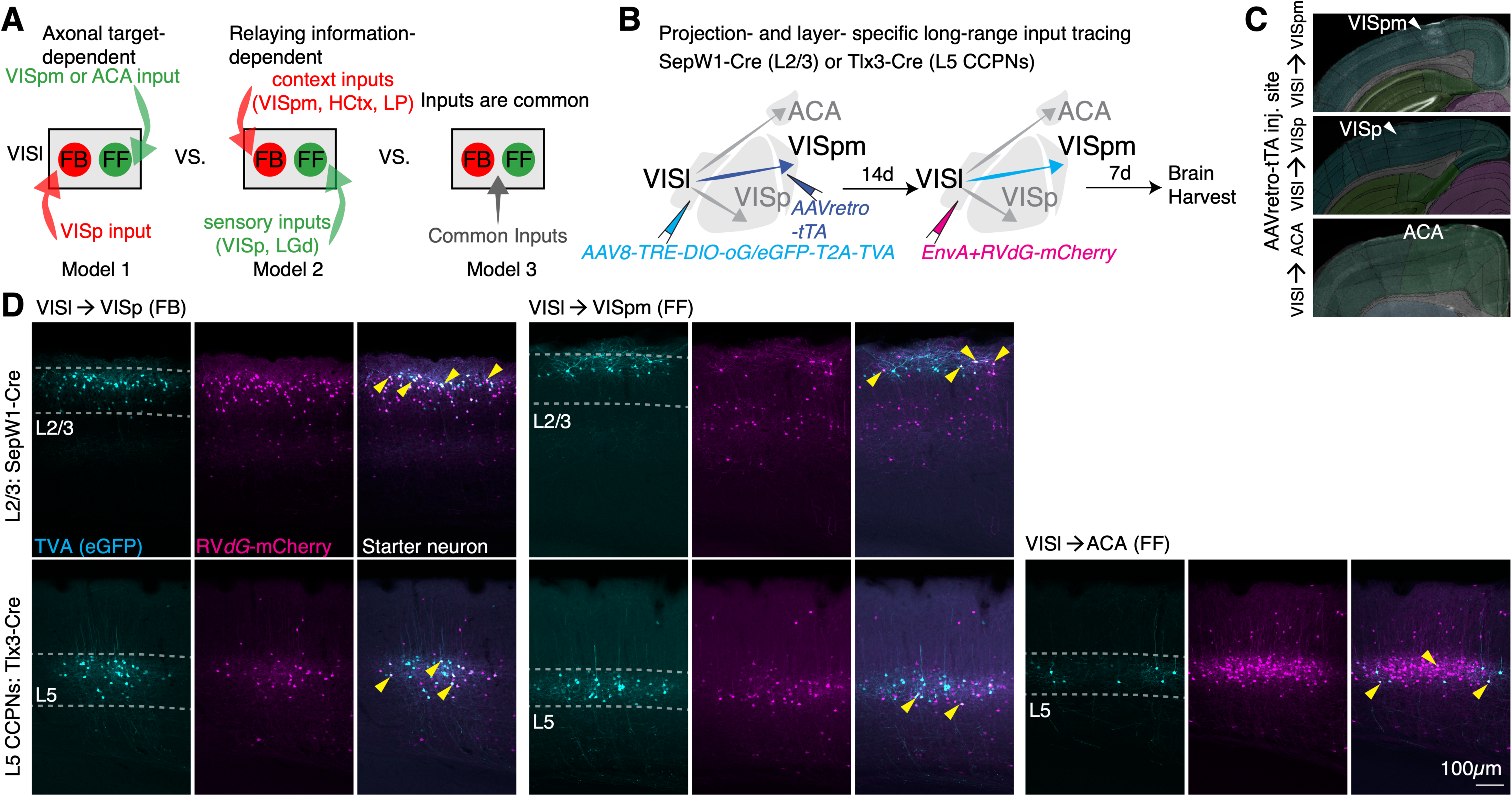
Long-distance connectivity models and projection-and layer-specific rabies tracing strategy. **(A)** Three proposed models of long-distance monosynaptic input connectivity to feedforward (FF) and feedback (FB) neurons. **(B)** Schematic of the intersectional rabies tracing strategy for projection-and layer-specific long-distance input tracing to VISl neurons. **(C)** Representative scanning microscope images overlaid with the Allen Brain Common Coordinate Framework reference atlas (Wang et al., 2020), showing AAVretro-tTA targeting in three different areas: VISpm, VISp, and ACA. **(D)** Representative confocal microscope images showing eGFP+mCherry+ starter neurons (arrowheads) in VISl projecting to VISp, VISpm, and ACA in *SepW1-Cre* or *Tlx3-Cre* mice.

To differentiate between these three models, we mapped and characterized the brain-wide long-distance inputs to distinct feedforward and feedback CCPNs in VISl. We implemented an intersectional rabies tracing strategy called cTRIO to map monosynaptic inputs in a projection-and layer-specific manner. We first injected a helper virus mixture of AAV8-TRE-DIO-oG (rabies glycoprotein) (Kim et al., 2016) and AAV8-TRE-DIO-eGFP-T2A-TVA into VISl, along with AAVretro-nEF-lox66/71-tTA (hereafter AAVretro-tTA) into projection targets, including VISpm, VISp, and ACA, in L2/3-specific *SepW1-Cre* or L5 CCPN-specific *Tlx3-Cre* transgenic mice (Gerfen et al., 2013). This ensured Cre-and tTA-dependent expression of the avian TVA receptor and rabies glycoprotein (oG). This approach enabled selective labeling of feedforward and feedback neurons in L2/3 or L5 (Fig. 2D). 14 days later, EnvA-pseudotyped glycoprotein (G)-deleted rabies virus (EnvA+RVdG)-mCherry was injected into the same VISl location to enable monosynaptic input tracing from the starter neurons. After seven days, brains were harvested, sectioned, immunostained with goat anti-eGFP (Rockland 600-101-215, RRID:AB_218182) and rabbit anti-dsRed (Clontech 632496, RRID:AB_10013483) antibodies, and imaged to quantify brain-wide long-distance inputs.

### Control experiments to optimize layer-and projection-specific monosynaptic rabies tracing

We conducted a series of experiments to optimize our monosynaptic rabies tracing approach. Our goal was to minimize false-positive long-distance labeling and prevent contamination of the starter neuron population. Such issues could arise from ‘leaky’ expression of transgenes such as TVA and oG independent of Cre and tTA activity (Wall et al., 2010; Lavin et al., 2020), as well as from unintended damage to long-distance axonal tracts in the white matter (Watakabe and Hirokawa, 2018) (Fig. 3).

**Figure 3.**
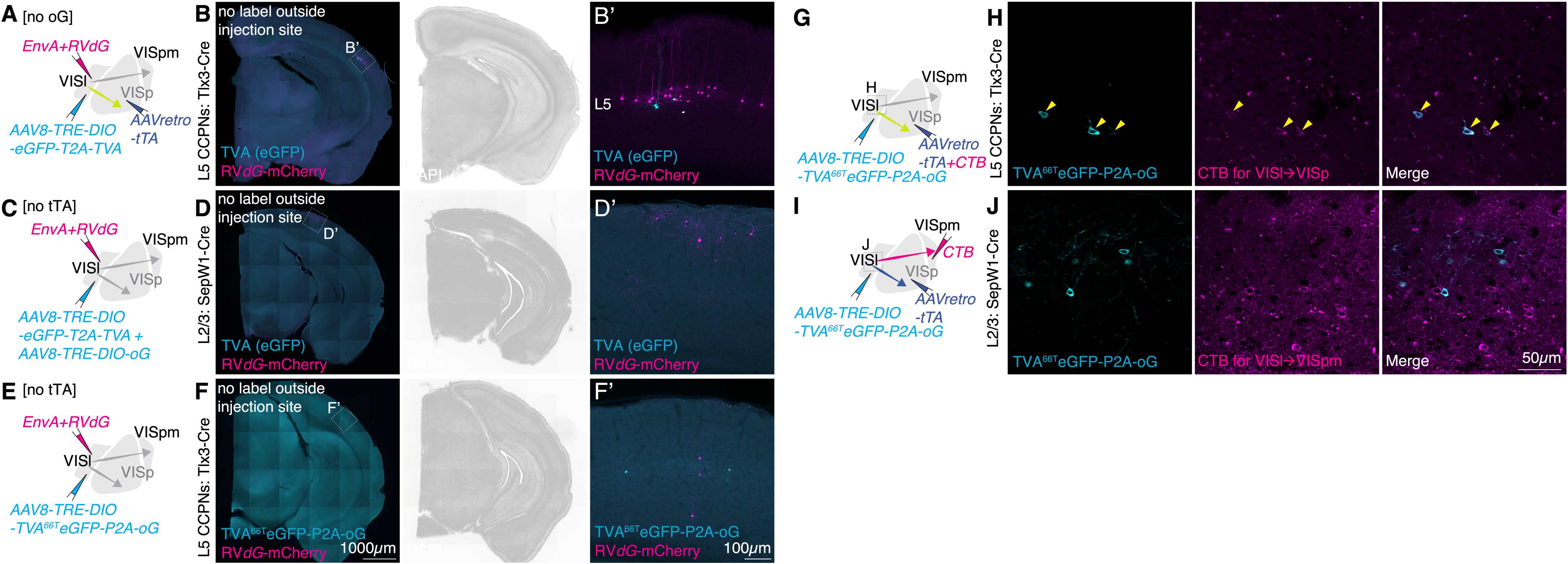
Control brains. **(A)** Schematic of the viral genetic tracing strategy for control injections without oG (rabies glycoprotein). **(B)** Fluorescence microscope images showing the absence of labeled cells outside the injection sites **(B’)**. **(C,E)** Schematic of the viral genetic tracing strategy for control injections without tTA. **(D-D’, F-F’)** Fluorescence microscope images showing the absence of labeled cells outside the injection sites **(D’, F’)**. **(G, I)** Schematic of the viral genetic tracing strategy used to verify the axonal projection targets of labeled starter neurons (TVA^66T^eGFP-P2A-oG+) by CTB injections into either the expected projection target, VISp **(G-H)**, or a non-target area, VISpm **(I-J)**. **(H)** Yellow arrowheads indicate starter neurons co-labeled with eGFP (cyan) and CTB (magenta), indicating their axonal projections to VISp.

**Figure 4.**
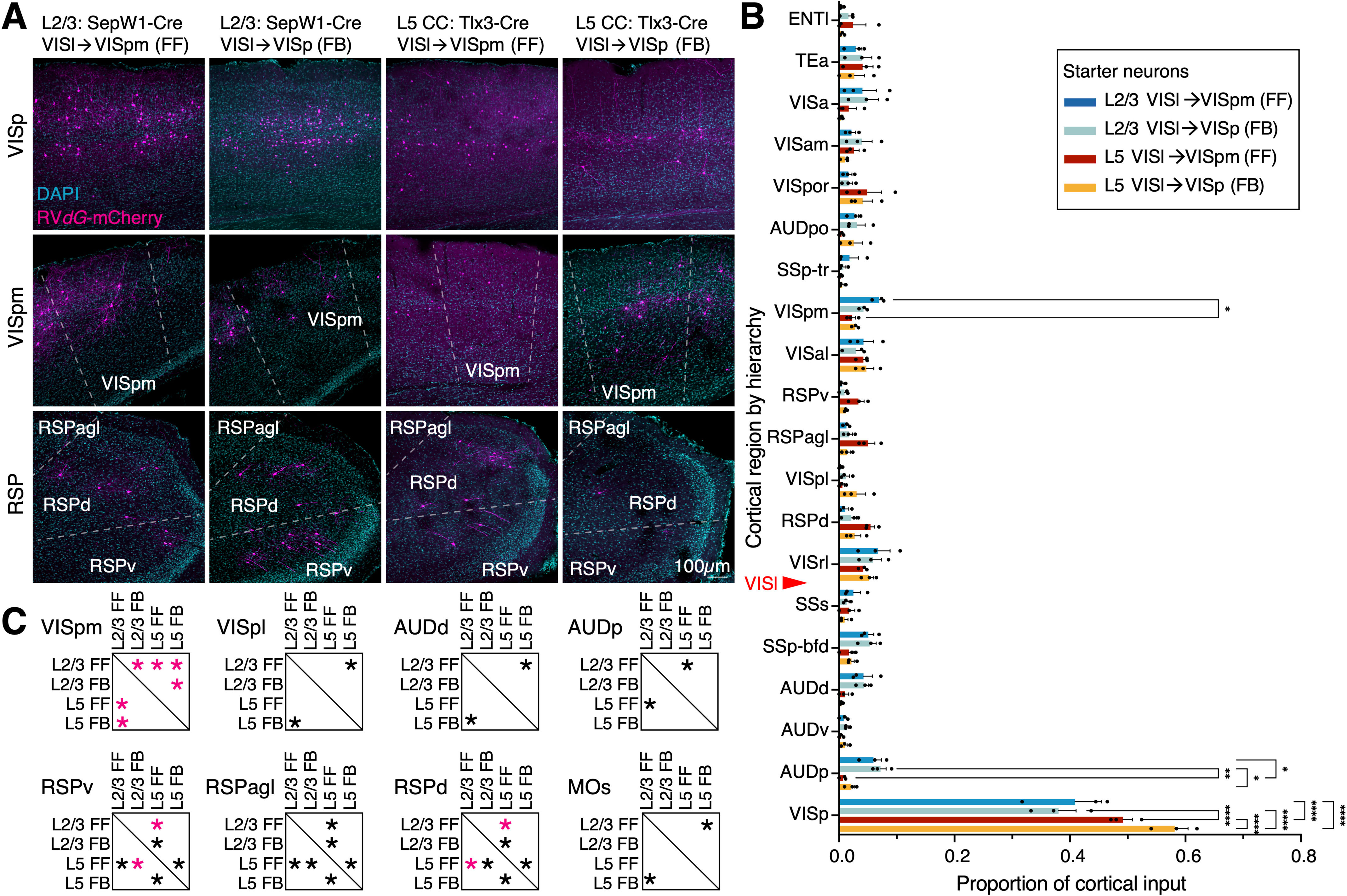
Long-distance input mapping of layer-and projection-specific CCPNs in VISl. **(A)** Representative confocal images showing long-distance mCherry+ input neurons in LP, VISp, VISpm, and RSP for four distinct feedforward and feedback CCPN types in VISl of *SepW1-Cre* or *Tlx3-Cre* mice. **(B)** The ipsilateral cortical proportion of input neurons across 20 cortical areas, sorted by cortical hierarchy (Harris et al., 2019). The red arrowhead indicates the rank of VISl along the cortical hierarchy. n = 3 mice per each group. Two-way ANOVA with Tukey’s multiple comparisons test, * p<0.05, ** p<0.01, *** p<0.001, **** p<0.0001. **(C)** Results of one-way ANOVA with Fisher’s Least Significant Difference (LSD) post-hoc test for eight selected input regions. *(black) p<0.05, *(magenta) p<0.01. Data are represented as mean ± SEM.

To examine the absence of false-positive long-distance rabies labeling, we first injected AAV8-TRE-DIO-eGFP-T2A-TVA (without oG) into VISl and AAVretro-tTA into VISp of *Tlx3-Cre* mice (Fig. 3A). 14 days later, we administered EnvA+RVdG-mCherry into VISl and harvested the brains seven days post-injection (Fig. 3A). As expected, in the absence of oG, no mCherry+ rabies-labeled neurons were detected outside of the injection site (VISl) (Fig. 3B-B’).

In additional controls, we injected either a mixture of AAV8-TRE-DIO-eGFP-T2A-TVA and AAV8-TRE-DIO-oG (Fig. 3C), or AAV8-TRE-DIO-TVA^66T^eGFP-P2A-oG alone (Fig. 3E) into VISl or VISp, but without AAVretro-tTA injections. In both cases, following EnvA+RVdG-mCherry and a seven-day incubation period, no mCherry+ rabies-labeled neurons outside of the injection site (Fig. 3C-D’, E-F’)

Next, to assess the purity of starter neurons and verify whether L2/3 or L5 VISl→VISp CCPNs contained only VISl→VISp neurons, we injected AAV8-TRE-DIO-TVA^66T^eGFP-P2A-oG into VISl and a mixture of AAVretro-tTA and Alexa Fluor 647-conjugated cholera toxin subunit B (CTB) into VISp of *Tlx3-Cre* mice (Fig. 3G-H), and collected brains three weeks later. Confocal imaging showed that TVA^66T^eGFP-P2A-oG labeled L5 neurons in VISl were also labeled with CTB, showing that they were VISl→VISp neurons (Fig. 3H).

To determine whether VISl→VISpm neurons were included in the L2/3 VISl→VISp starter population due to passing axons in the grey matter of lower cortical layers (Watakabe and Hirokawa, 2018), we injected AAV8-TRE-DIO-TVA^66T^eGFP-P2A-oG into VISl, AAVretro-tTA into VISp, and CTB into VISpm of *SepW1-Cre* mice (Fig. 3I-J). Confocal images of brain sections harvested three weeks later showed that TVA^66T^eGFP-P2A-oG expressing neurons were not labeled with CTB, indicating that virus-labeled starter neurons did not include VISl→VISpm neurons (Fig. 3J). These experiments validate our approach, ensuring specificity in identifying monosynaptic inputs while preventing false positives and potential starter population contamination.

### Long-distance cortical inputs onto four different CCPN types in VISl

We examined whether the four types of feedforward and feedback CCPNs (L2/3 VISl→VISpm, L2/3 VISl→VISp, L5 VISl→VISpm, L5 VISl→VISp) in VISl receive similar or distinct monosynaptic inputs from other cortical areas within the ipsilateral hemisphere (Fig. 4 and Table 2). Rabies-labeled long-distance input neurons to those four types of CCPNs were distributed throughout both visual and non-visual cortical areas (Fig. 4A), with the highest proportion originating from VISp (L2/3 VISl→VISpm 0.409 ± 0.046, L2/3 VISl→VISp 0.380 ± 0.030, L5 L5 VISl→VISpm 0.492 ± 0.016, L5 VISl→VISp 0.582 ± 0.023, Fig. 4B). We quantified and ranked the top 20 cortical regions contributing inputs along the cortical hierarchy (Harris et al., 2019). While overall input patterns among the four CCPN types were largely similar, we identified key differences. L5 VISl→VISp feedback neurons received a greater proportion of long-distance inputs from VISp compared to L5 VISl→VISpm feedforward neurons (Fig. 4B).

**Table 2.**
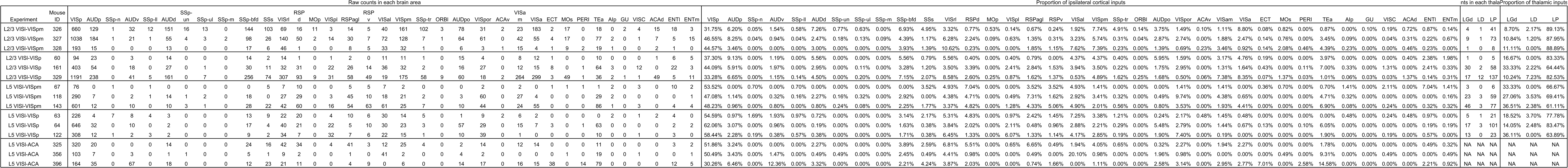
Quantification of rabies tracing in individual mouse brains.

Similarly, L2/3 VISl→VISpm feedforward neurons received a significantly greater proportion of their inputs from VISpm than L2/3 VISl→VISp feedback neurons (Fig. 4C).

Further statistical analysis using one-way ANOVA revealed that at least one of the four CCPN types exhibited a significantly different proportional input than other CCPNs in eight out of the 20 cortical areas analyzed (Fig. 4C): VISpm, VISpl, AUDd, AUDp, RSPv, RSPagl, RSPd, and MOs. Overall, these findings suggest that while the four CCPN types receive broadly comparable long-distance cortical inputs, they exhibit distinct preferences for specific cortical regions, reflecting their functional roles in cortical processing. Notably, L5 VISl→VISpm CCPNs receive proportionally more retrosplenial cortex input, suggesting a greater role in visuo-spatial memory, and L2/3 CCPNs regardless of projection receive proportionally more auditory cortex input suggesting a greater role for L2/3 CCPNs in audio-visual integration (Fig. 4B-C).

We next investigated the laminar distribution of long-distance presynaptic inputs from VISp and VISpm to four distinct types of feedforward and feedback CCPNs in VISl. Although prior studies have reported preferential long-distance connectivity between superficial layers and between deep layers (DeNardo et al., 2015; Kim et al., 2015), it remains unclear whether CCPNs in specific layers of a given cortical area receive layer-specific long-distance inputs, depending on their feedforward or feedback projections. To address this, we quantified the proportion of input neurons across each layer of VISp and VISpm (Fig. 5A-B) for four distinct CCPN types in VISl: L2/3 VISl→VISpm, L2/3 VISl→VISp, L5 VISl→VISpm, and L5 VISl→VISp. With a few exceptions, the laminar distribution of input neurons in VISp and VISpm did not reveal statistically significant differences across these neuronal groups (Fig. 5A-B).

**Figure 5.**
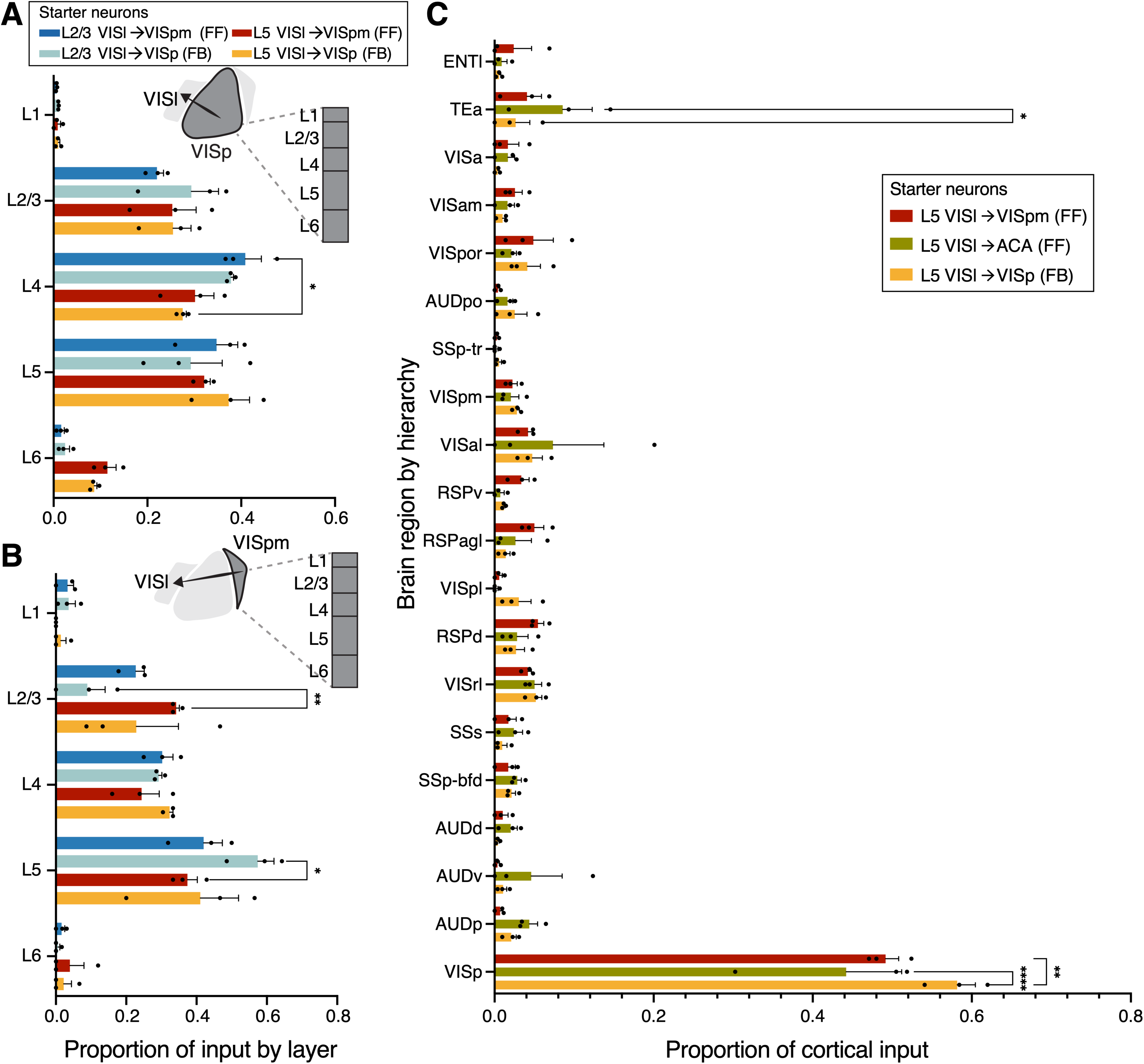
Further analyses of long-distance input neurons to CCPNs in VISl. **(A)** The distributions of presynaptic input neurons across VISp layers to four distinct types of CCPNs in VISl. **(B)** The distribution of presynaptic input neurons across VISpm layers to four distinct types of CCPNs in VISl. **(C)** The proportion of long-distance input neurons located in each of the 20 cortical regions for the three L5 CCPN types in VISl. n = 3 mice per each group for each figure. Two-way ANOVA with Tukey’s multiple comparisons test, * p<0.05, ** p<0.01, **** p<0.0001. Data are represented as mean ± SEM.

We investigated whether feedforward neurons in VISl that project to cortical areas with higher hierarchy scores than VISpm exhibit more distinct input connectivity patterns from those of VISl→VISp compared to VISl→VISpm neurons. Previous studies have shown that the cortical hierarchy in rodents is relatively shallow compared to that of primates (Harris et al., 2019; D’Souza et al., 2022). The hierarchy scores based on layer-termination patterns of cortico-cortical connection (Harris et al., 2019) for select cortical areas are as follows: VISp:-0.239, VISl:-0.148, VISpm: 0.074, ACAd:-0.060, ACAv: 0.404, and MO: 0.282., To determine if the shallow hierarchy in the mouse visual cortex was masking hierarchy dependent input differences, we injected AAVretro-tTA into ACA/MOs and a mixture of AAV8-TRE-DIO-oG and AAV8-TRE-DIO-eGFP-T2A-TVA into VISl of *Tlx3-Cre* mice to label VISl→ACA/MOs (hereafter, VISl→ACA) neurons and traced their monosynaptic inputs across the cortex. We then compared these results with monosynaptic input tracing of L5 VISl→VISpm feedforward neurons and L5 VISl→VISp feedback neurons (Fig. 5C). We found broadly similar input connectivity profiles across these three L5 CCPN subtypes. Thus, the input connectivity of L5 VISl→ACA CCPNs is comparable to that of VISl→VISpm CCPNs and does not exhibit greater differences from VISl→VISp CCPNs than from VISl→VISpm CCPNs. Notably, the proportion of inputs from VISp to VISl→VISp neurons was higher than that to VISl→VISpm and VISl→ACA neurons (Fig. 5C).

### Long-distance thalamic input neuron profiles onto four different CCPN types in VISl

Next, we examined whether the four types of feedforward and feedback CCPNs in VISl receive distinct presynaptic inputs from visual thalamic regions. In the mouse visual system, LGd projects to VISl and other higher visual cortical areas in addition to VISp, while LP and the laterodorsal nucleus of the thalamus (LD) also provide input to VISl and other visual cortical regions (Bienkowski et al., 2019).

In rabies-traced brain sections, we identified three thalamic nuclei, LGd, LD, and LP, and quantified the proportion of rabies-labeled neurons among all these input areas (Fig. 6A). Across all four CCPN types, LP provided the majority of thalamic inputs, followed by LGd and LD, consistent with a previous study (Yao et al., 2023a) (Fig. 6B). While the overall proportions of thalamic inputs were similar, we observed that L2/3 VISl→VISpm neurons received a lower proportion of inputs from LGd compared to L5 VISl→VISpm neurons (L2/3 VISl→VISpm: 0.102 ± 0.008 and L5 VISl→VISpm 0.323 ± 0.027, p=0.0030, two-way ANOVA with Tukey’s multiple comparisons test) (Fig. 6B). Conversely, L2/3 VISl→VISpm neurons received a higher proportion of inputs from LP than L5 VISl→VISpm neurons (L2/3 VISl→VISpm 0.887 ± 0.004, L5 VISl→VISpm 0.657 ± 0.024, p=0.0021, two-way ANOVA with Tukey’s multiple comparisons test) (Fig. 6B).

**Figure 6.**
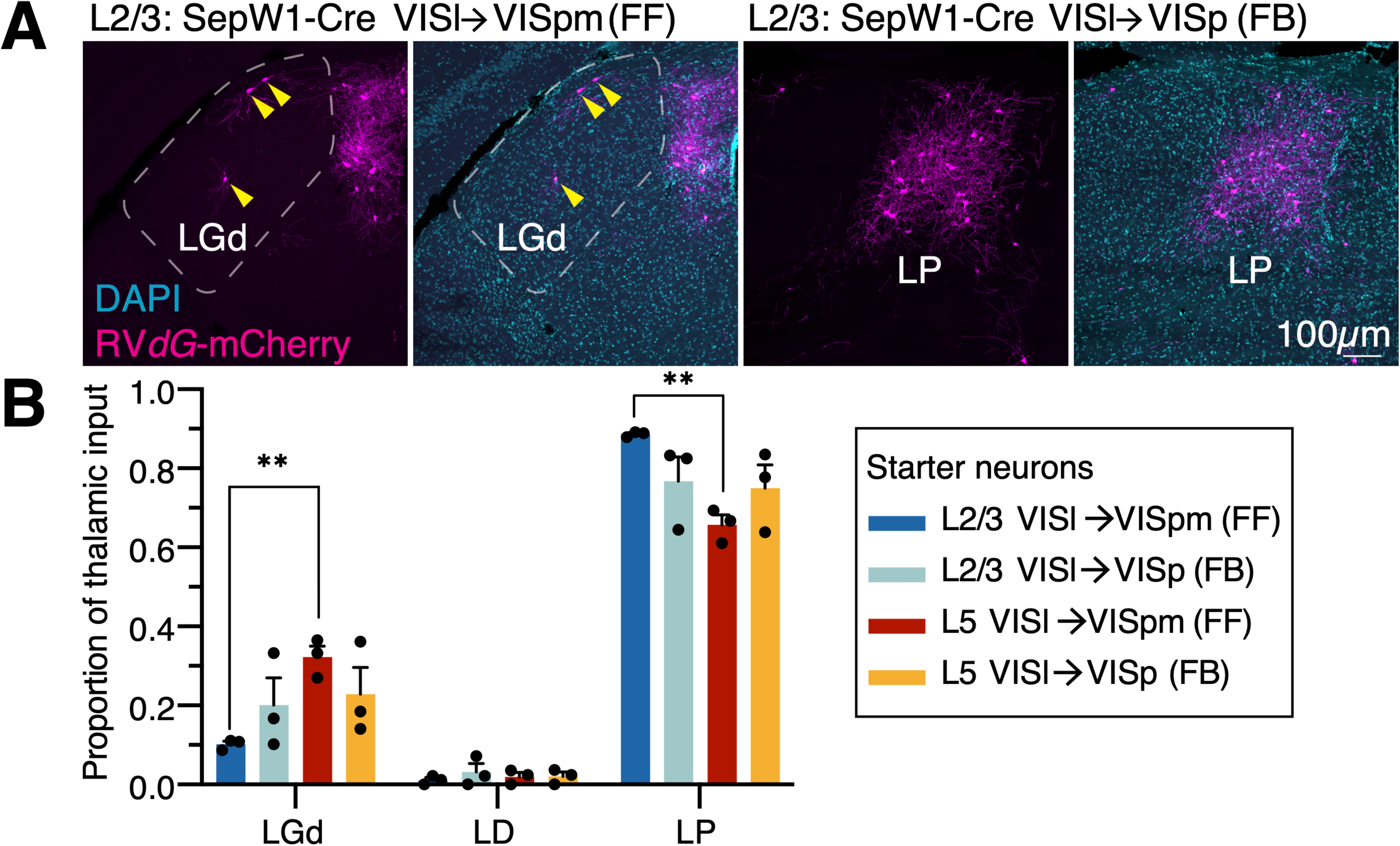
Long-distance thalamic input mapping of layer-and projection-specific CCPNs in VISl. **(A)** Representative confocal microscope images showing rabies-labeled mCherry+ input neurons in LGd or LP. **(B)** Proportions of long-distance inputs from each visual thalamic region to four distinct types of feedforward and feedback starter neurons in VISl. n = 3 mice per each group. Two-way ANOVA with Tukey’s multiple comparisons test, * p<0.05. Data are represented as mean ± SEM.

## DISCUSSION

Using a layer-and projection-specific monosynaptic rabies tracing approach, we mapped brain-wide inputs to five distinct types of feedforward and feedback CCPNs in VISl. This study provides the first comprehensive connectivity maps to investigate the organizational principles governing feedforward and feedback neurons in relation to their laminar positions and the hierarchical positions of their target areas. Our findings indicate that long distance input connectivity patterns are largely comparable across CCPN subtypes, supporting Model 3 (Fig. 2A), but notable differences emerge based on their laminar and projection identities. First, L2/3 VISl→VISpm CCPNs received a higher proportion of inputs from their own projection target, VISpm, compared to L2/3 VISl→VISp and other L5 CCPNs. Similarly, L5 VISl→VISp CCPNs received a higher proportion of inputs from VISp, their projection target, compared to L5 VISl→VISpm or VISl→ACA CCPNs. These results are in accordance with Model 1 (Fig. 2A), in which monosynaptic inputs preferentially originate from the neurons’ axonal targets (Kim et al., 2020; Siu et al., 2021). These insights provide a foundation for future functional and theoretical studies aimed at understanding brain-wide cortical computations at the cellular level.

The input-output organization of neural networks provides crucial insights into the information they receive and the signals they transmit for functional processing. Therefore, understanding neuronal connectivity patterns is essential for a comprehensive understanding of neural circuit function and computation (Zeng and Sanes, 2017). Previous studies have demonstrated that distinct neuronal cell types in the mammalian brain exhibit characteristic input connectivity patterns. Even within broader neuronal classes, spatially intermingled subtypes display subtype-specific input connectivity (e.g., dopaminergic D1 receptor versus D2 receptor expressing spiny projection neurons in the striatum or serotonergic neuron subtypes in the dorsal raphe; see also whole-brain rabies tracing data) (Wall et al., 2016; Ren et al., 2018). A recent study found that functionally distinct neuronal types in layer 2/3 of the mouse somatosensory cortex exhibit distinct brain-wide input connectivity patterns (Inácio et al., 2025). For cortico-cortical connectivity in the visual cortices, a previous study found that VISp CCPNs projecting to VISpm or VISal received inputs from the same respective areas, suggesting a like-to-like connectivity pattern (Kim et al., 2020). A previous study observed laminar differences in like-to-like connectivity using Channelrhodopsin-2-assisted circuit mapping, where there is stronger like-to-like connectivity in L5 compared to L2/3 neurons (Young et al., 2021). This observation of like-to-like connectivity is further supported by a study in macaques, where V1 neurons projecting to V2 receive monosynaptic feedback inputs from V2 but not from other V1-projecting areas (Siu et al., 2021). In the mouse VISl, the secondary visual cortex, projection-specific input connectivity is observed, though it appears less pronounced than in CCPNs of VISp, the primary visual cortex. One possible explanation is that VISl→VISpm and VISl→VISp neurons share similar medial axonal projection trajectories, whereas VISp→VISpm and VISp→VISal (VISp CCPNs projecting to anterolateral, AL, visual cortex) neurons exhibit medial versus lateral projection patterns. Another, not mutually exclusive, possibility is that the cortical hierarchy in the mouse visual system is relatively shallow, making it more challenging to detect strong hierarchical connectivity features.

Although little is known about LP and LGd projections to VISl, our observed LP:LGd input ratios are largely consistent with previous rabies-tracing data examining thalamic inputs to layer-specific VISl populations (Yao et al., 2023a). Notably, we found that VISl L5 FF neurons projecting to VISpm receive proportionally more LGd input compared to their L2/3 counterparts, suggesting a Model 2 input organization in which FF neurons receive stronger input from feedback-associated areas such as LGd. However, this pattern is not observed in L2/3 FF neurons, indicating a non-symmetrical input organization between FF and FB neurons across layers. Importantly, our data describe input proportions rather than functional strengths; future quantitative and functional studies, such as electrophysiological recordings or other synaptic connectivity measures, will be necessary to clarify the strength and dynamics of these thalamocortical connections.

To validate the projection identities of our starter neurons, we performed CTB injections into both target and non-target projection areas (Fig. 3G-J). However, given the collateralization properties of some CCPNs (Han et al., 2018; Peng et al., 2021), it is possible that a subset of starter neurons also projects to additional areas beyond the intended targets. Notably, axonal reconstructions and RNA barcode-based mapping (MAPseq) by Han et al. demonstrated that approximately 88% of VISp CCPNs project to only one or two of the six higher visual areas. Similarly, Berezovskii et al. (2011) using dextran amine tracers reported that FF (to VISp) and FB (to VISal) CCPNs in VISl are largely non-overlapping, with less than 5% overlap. Our observations of VISl axonal collateralization are consistent with these findings, while acknowledging potential differences between VISl and VISp as well as variations in experimental approaches. Nonetheless, we note that potential collateral projections of VISl CCPNs to areas beyond ACA, VISp, and VISpm could introduce variability in rabies tracing results and complicate efforts to restrict analyses exclusively to single projection-defined populations.

Monosynaptic rabies tracing is a powerful approach for mapping anatomical connections across the brain; however, this method does not provide information about synaptic strength or other functional properties of these connections (Callaway and Luo, 2015). For example, although feedforward and feedback CCPNs may receive similar proportions of inputs from a given area, it remains unclear whether these input neurons are functionally distinct—for example, whether they exhibit different receptive field properties or belong to different functional subtypes. To gain additional insight into the functional differences and similarities between feedforward and feedback CCPNs, future studies could complement rabies tracing with electrophysiological or imaging approaches. Additionally, advances in high-throughput molecular profiling, including single-cell RNA sequencing, have revealed that cortical neurons within a single layer can be further classified into distinct subtypes based on unsupervised clustering (Tasic et al., 2018; Yao et al., 2023b). Beyond their broad excitatory versus inhibitory or laminar identities, it would be valuable to examine whether long-distance input neurons exhibit specific connectivity patterns among these molecularly defined subtypes of CCPNs (Condylis et al., 2022).

It is also possible that, despite receiving similar long-distance inputs, feedforward and feedback neurons differ in their local circuit architecture. To further test Model 3 (Fig. 2A) and uncover additional anatomical distinctions, future studies using TVA^66T^-dependent rabies tracing for local inputs, combined with molecular characterization of input neurons and electrophysiological profiling would be informative.

In this study, we used feedforward (to VISpm or ACA) and feedback (to VISp) CCPNs in VISl as a model to investigate connectivity differences. Given that mammalian visual cortical areas are organized into dorsal and ventral streams with strong intra-stream connectivity, mapping inputs to feedforward and feedback neurons targeting areas within a stream, such as POR, a higher-order ventral stream area, could offer valuable insights <u>(D’Souza et al. 2022; Harris et al.</u> <u>2019)</u>. Although POR represents an informative target for examining feedforward and feedback connectivity, precise stereotaxic targeting remains technically challenging. Future studies including POR could further clarify ventral stream-specific connectivity. Another key question for future research is whether similar connectivity profiles would be observed in other cortical regions, such as secondary somatosensory cortex projecting to primary somatosensory cortex as feedback CCPNs and secondary somatosensory cortex projecting to primary and secondary motor cortices as feedforward CCPNs (Yamashita et al., 2018). Given that the cortical hierarchy in mice is relatively shallow compared to that of higher mammals (Harris et al., 2019; D’Souza et al., 2022), it would also be interesting to explore these questions in species with a more robust hierarchical organization, such as marmosets or macaques.

## Author Contributions

Richard G. Dickson performed investigations, supervised students, collected and curated data, analyzed data, prepared visualizations, and contributed to writing the original draft and revisions. Matthew W. Jacobs performed investigations, supervised students, and collected data. John M.

Ratliff performed investigations, collected and analyzed data, and contributed to writing and revisions. Alec L. R. Soronow performed investigations. Faye An and Walid A. Yuqob collected and analyzed data. Euiseok J. Kim conceived and designed the study, supervised the project, analyzed and visualized data, contributed resources, acquired funding, and contributed to writing the original draft and revisions. All authors had full access to the data, take responsibility for its integrity, and approved the final manuscript.

## Abbreviations

AAV: Adeno associated virus
ACA: Anterior cingulate area
ACAd: Anterior cingulate area dorsal
ACAv: Anterior cingulate area ventral
Alp: Agranular insular area, posterior part
AUDd: Dorsal auditory area
AUDp: Primary auditory area
AUDpo: Posterior auditory area
AUDv: Ventral auditory area
CCPN: Cortico-cortical projection neurons
CTB: Cholera toxin subunit B
cTRIO: Cell specific tracing the relationship between input and output
ECT: Ectorhinal area
ENTl: Entorhinal area, lateral part
ENTm: Entorhinal area, medial part, dorsal zone
EnvA: Envelope protein of avian leukosis virus
FB: feedback neurons
FF: feedforward neurons
GU: Gustatory areas
HVA: Higher visual area
LD: Lateral dorsal nucleus of thalamus
LGd: Lateral geniculate nucleus dorsal
LP: Lateral posterior nucleus of the thalamus
Mop: Primary motor area
Mos: Secondary motor cortex
oG: optimized rabies glycoprotein
ORBl: Orbital area, lateral part
PBS: Phosphate buffered saline
PERI: Perirhinal area
PFA: paraformaldehyde
RSP: Retrosplenial cortex
RSPagl: Retrosplenial cortex agranular
RSPd: Retrosplenial cortex dorsal
RSPv: Retrosplenial cortex ventral
SSp-bfd: Primary somatosensory area, barrel field
SSp-ll: Primary somatosensory area, lower limb
SSp-m: Primary somatosensory area, mouth
SSp-n: Primary somatosensory area, nose
SSp-ul: Primary somatosensory area, upper limb
SSp-un: Primary somatosensory area, unassigned
SSp-tr: Primary somatosensory area, trunk
SSs: Supplemental somatosensory area
TEa: Temporal association area
tTa: Tetracycline trans activator
TVA: Receptor for avian leukosis virus
VISa: Anterior area secondary visual cortex
VISal: Anterolateral secondary visual cortex
VISam: Anteromedial secondary visual cortex
VISC: Visceral area
VISl (LM): Lateral secondary visual cortex
VISli: Latero intermediate secondary visual cortex
VISp: Primary visual cortex
VISpl: Posterolateral secondary visual cortex
VISpm: Posteromedial secondary visual cortex
VISpor: Postrhinal secondary visual cortex
VISrl: Rostrolateral secondary visual cortex.

---

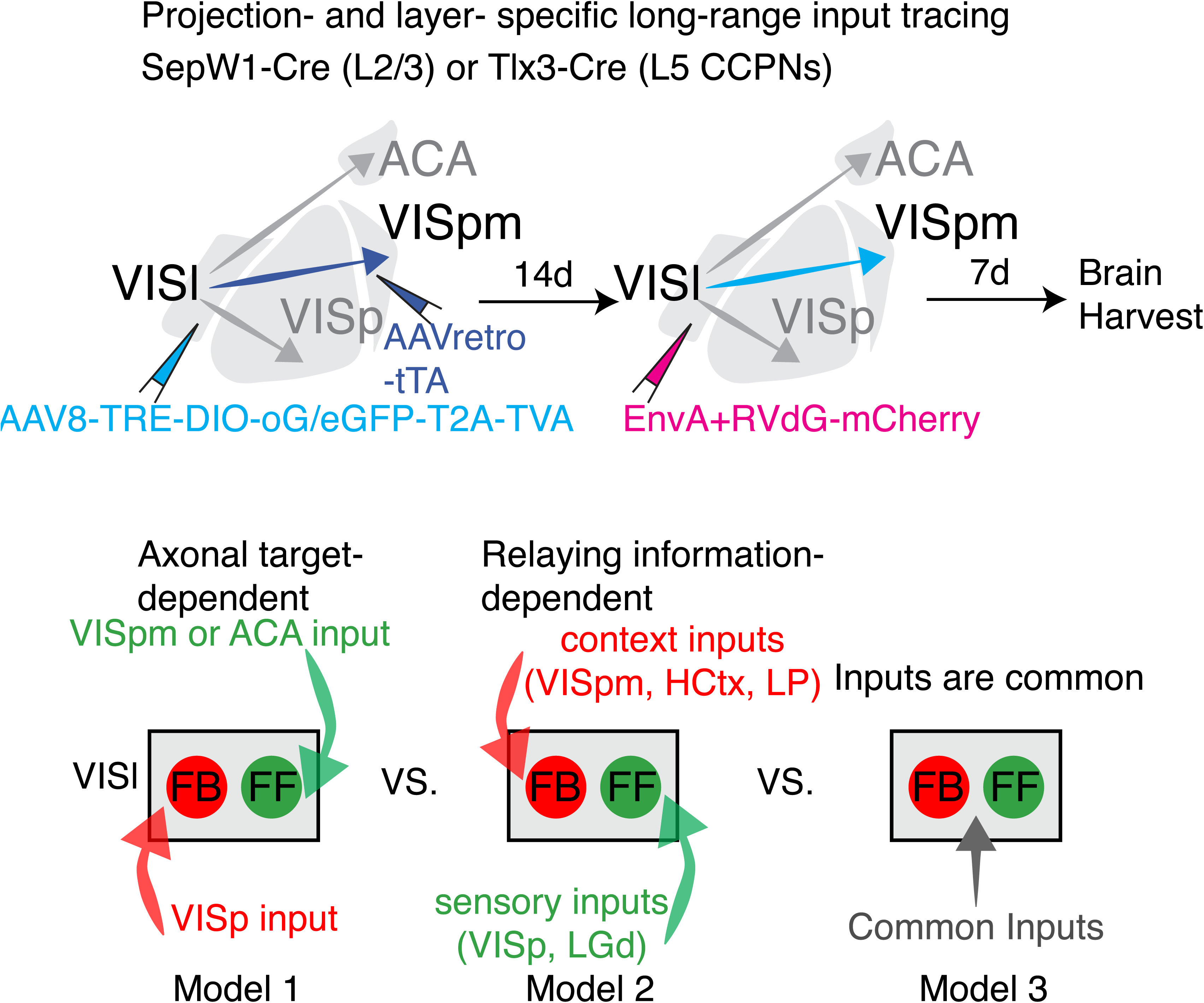

